# A hyperbolic topological atlas reveals polyamine steering of a shared developmental manifold in Arabidopsis

**DOI:** 10.64898/2026.06.23.733990

**Authors:** Jan Zdražil, Lingping Kong, Emmanuel Flores-Hernández, Margarita Rodríguez-Kessler, Pavel Klimeš, Lukáš Spíchal, Nuria De Diego, Václav Snášel

**Author notes:** **Corresponding authors:** Nuria De Diego, Václav Snášel.

## Abstract

High-throughput plant phenotyping captures development at scale, yet image-rich screens are still often reduced to static trait summaries. We tested whether nutrient availability, polyamine priming, concentration, and their transport reshape Arabidopsis rosette development by generating distinct morphologies or by changing residence along a common trajectory. We analyzed 138,223 time-resolved rosette images from Col-0 and five mutants involved in polyamine transport (*put1-5*) primed to putrescine, spermidine, spermine, dose, and nutrient regimes using a self-supervised vision backbone, Poincaré embedding, hyperbolic Mapper, and manifold straightening. The data form a single connected developmental manifold with 410 nodes and 746 edges, organized from an early, low-nutrient-biased hub through high-betweenness transition corridors to two late, nutrient-enriched terminal regions. Polyamine identity stratifies this manifold by developmental phase: putrescine enriches early states, spermidine occupies transition corridors, and spermine marks late compact rosettes. Nutrient richness and dose change distal occupancy, whereas put genotypes alter dwell time within shared regions rather than producing separate topologies. Manifold straightening resolves these effects into a short early lateral deflection followed by convergence, yielding two scalar readouts, early transverse offset and distal occupancy, that summarize treatment action on a common morphodynamic scale. The framework converts large image screens into interpretable developmental geometry for image-based phenomics.

## Introduction

Plant morphogenesis is a continuous process in which organ initiation, expansion, overlap, and canopy closure translate genetic and environmental inputs into form. In rosette plants, high-throughput imaging captures these changes non-destructively, yet image time series are still often reduced to isolated descriptors such as projected area, color, compactness, or growth rate.[1–8] Recent AI-enabled phenomics studies have expanded what researchers can extract from large plant-image collections, but they also show that image acquisition alone is no longer the central challenge; the challenge is to convert image-rich screens into interpretable, reproducible developmental readouts.[4,5,9–11] Static traits remain valuable, but they do not directly answer a central developmental question: whether a perturbation creates a new morphogenetic route or changes the speed and residence of plants within states already available to the species.

Representation learning offers a way to recover this hidden continuity from images. Self-supervised vision models can encode complex plant morphology without manually defined traits, and manifold-learning guidance emphasizes the diagnostic interpretation of embeddings rather than their use as simple cluster maps.[4,5,9,12–15]. Ordinary low-dimensional projections, however, can flatten branch structure, compress transitional states, and encourage overinterpretation of clusters. A developmental atlas therefore needs both local sensitivity and global topology: it should preserve neighboring morphologies, retain long-range relationships, and allow experimental metadata to be projected only after the morphological space has been reconstructed.

Topological data analysis (TDA) and non-Euclidean representation learning provide that framework. Mapper graphs summarize high-dimensional data as connected regions and transitions; hyperbolic embeddings are well suited to hierarchies and branching progressions; and graph-based straightening can convert a curved manifold into interpretable pseudotime and transverse-offset coordinates.[16–20] These approaches are increasingly relevant to phenomics because development is not only a sequence of sizes, but a trajectory through a structured space of possible forms.

Polyamines provide a useful biological test case for geometry-aware analysis. Putrescine (Put), spermidine (Spd), and spermine (Spm) are small cationic metabolites associated with cell expansion, stress tolerance, flowering, and nutrient-dependent growth control.[21] Yet endpoint traits alone cannot resolve their effects on whole-rosette morphodynamics. The same projected area can arise from different canopy architectures, and a single treatment can change early expansion, transition timing, and terminal compactness in different ways.

Here we test whether nutrient status, polyamine priming, dose, and transport reorganize Arabidopsis rosette development by creating distinct morphological routes or by redistributing plants along a common developmental manifold. The dataset comprises 138,223 top-view rosette images spanning Col-0 and five *Atput(1-5)* transporter knockouts encoding each polyamine uptake transporter (*AtPUT*) gene family member, three polyamine priming regimes, non-zero dose levels plus unprimed controls, and full- versus low-MS nutrition. The workflow combines plant detection and super-resolution preprocessing with a DINOv2 feature backbone, Poincaré projection, hyperbolic space partitioning, Mapper graph reconstruction, logic-driven clustering, and manifold straightening (Figure 1).

**Figure 1.**
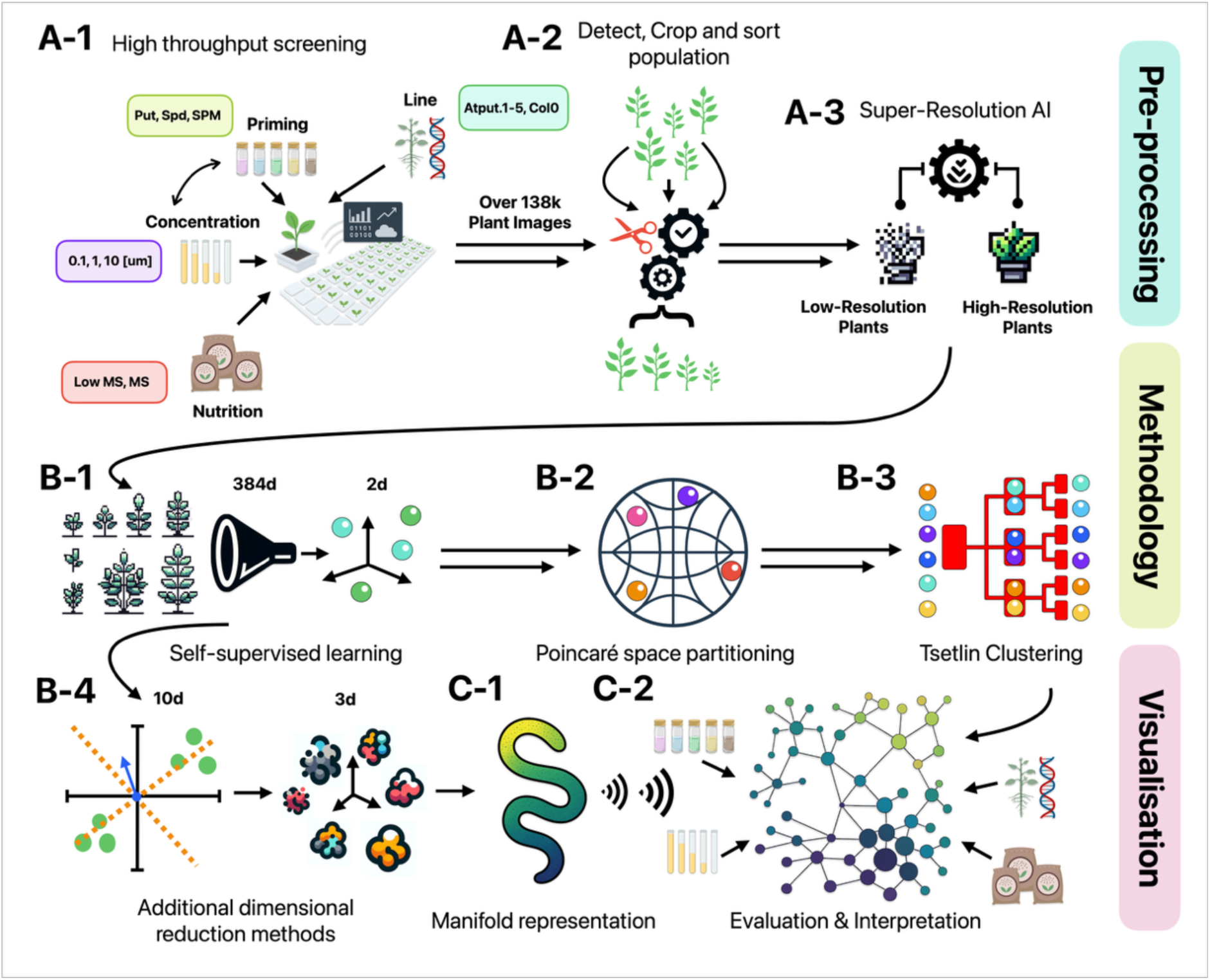
High-throughput screening and geometry-aware computational workflow. (A) Arabidopsis thaliana seedlings spanning polyamine priming (PUT, SPD, SPM), nutrient regimes, doses, and genotypes generate 138,223 plant images. (B) Automated preprocessing detects, crops, and enhances seedlings by super-resolution before self-supervised feature extraction, Poincaré embedding, and logic-based clustering. (C) Dimensionality reduction and manifold reconstruction convert raw phenotyping data into an interpretable developmental map linking time, treatment, genotype, and nutrition to emergent rosette morphology.

We separated geometry from annotation. Image features defined the manifold; we then used time, treatment, medium, concentration, and genotype to interpret occupancy, transitions, and deviations from the centerline. This design yielded two compact, biologically interpretable outputs: an early transverse offset that reports treatment-specific steering and a distal occupancy score that reports residence in late rosette programs. This morphodynamic scale allows complex phenotyping screens to be compared as trajectories rather than as disconnected collections of traits.

## Results

### A connected Mapper graph orders rosettes by developmental progression

Hyperbolic Mapper reconstruction generated a single connected graph of 410 nodes and 746 edges from 138,223 rosette images (Figure 2A-1). Coloring nodes by imaging round revealed a clear developmental gradient: early samples were concentrated in a compact hub, intermediate nodes formed narrow connector regions, and late samples occupied two distal arms. Representative rosette images embedded in the nodes supported this ordering, progressing from small, open seedlings to dense, overlapping canopies. Thus, the dominant structure recovered from images was not a set of isolated treatment clusters, but a connected developmental manifold with alternative late regions.

**Figure 2.**
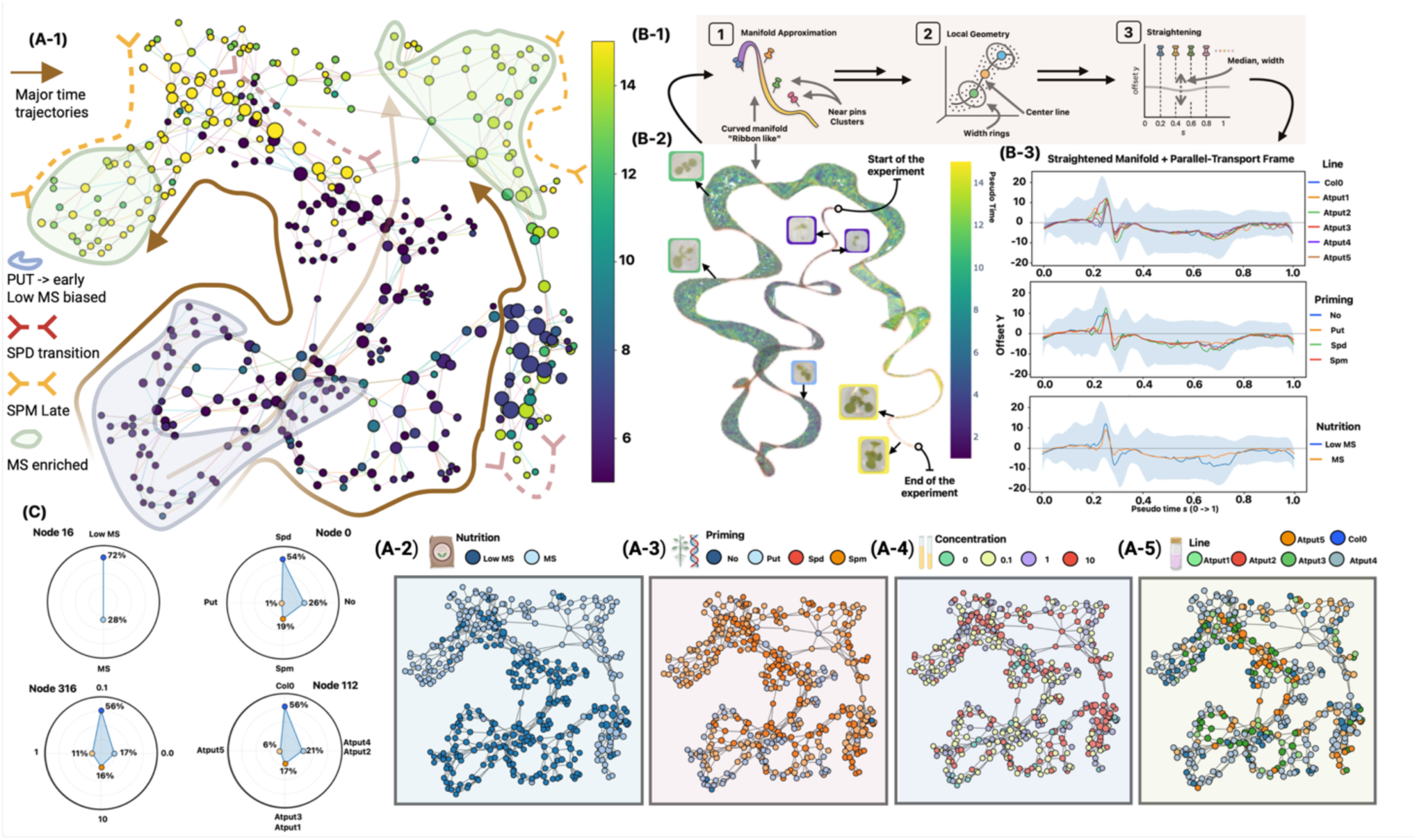
Shared rosette morphodynamic manifold and ribbon representation. (A-1) Mapper graph of 138,223 rosette images, organized from an early, low-MS-biased hub to two late, MS-enriched arms. Arrows denote major developmental directions; shaded regions mark Put-enriched early, Spd-enriched transition, and Spm-enriched late domains. (A-2 to A-5) Node-level overlays for nutrition, priming, concentration, and line show condition-specific occupancy on a shared topology rather than separate developmental paths. (B-1) Schematic of manifold straightening by medial-axis tracing and local coordinate-frame construction. (B-2) Three-dimensional representation of the curved ribbon derived from embeddings, colored by timepoint, with representative rosette images along the trajectory. (B-3) Parallel-transport ribbon coordinates quantify early lateral deflections and later convergence across line, priming, and nutrition. (C) Example node compositions highlight factor proportions.

### Polyamine identity maps onto developmental position

Projecting priming metadata onto the same graph revealed phase-specific enrichment without breaking graph connectivity (Figure 2A-3). The effect of putrescine (Put) was concentrated in the early hub, spermidine (Spd) was most frequent in connector nodes, and Spm was enriched at the distal arms, regardless of the used concentration. This distribution is consistent with a temporal partition of polyamine-associated morphology: Put marks early expansion states, Spd marks transitional states, and Spm marks compact late states. These data therefore indicate that polyamine identity changes where plants reside within a shared manifold rather than producing separate morphologies.

**Figure 3.**
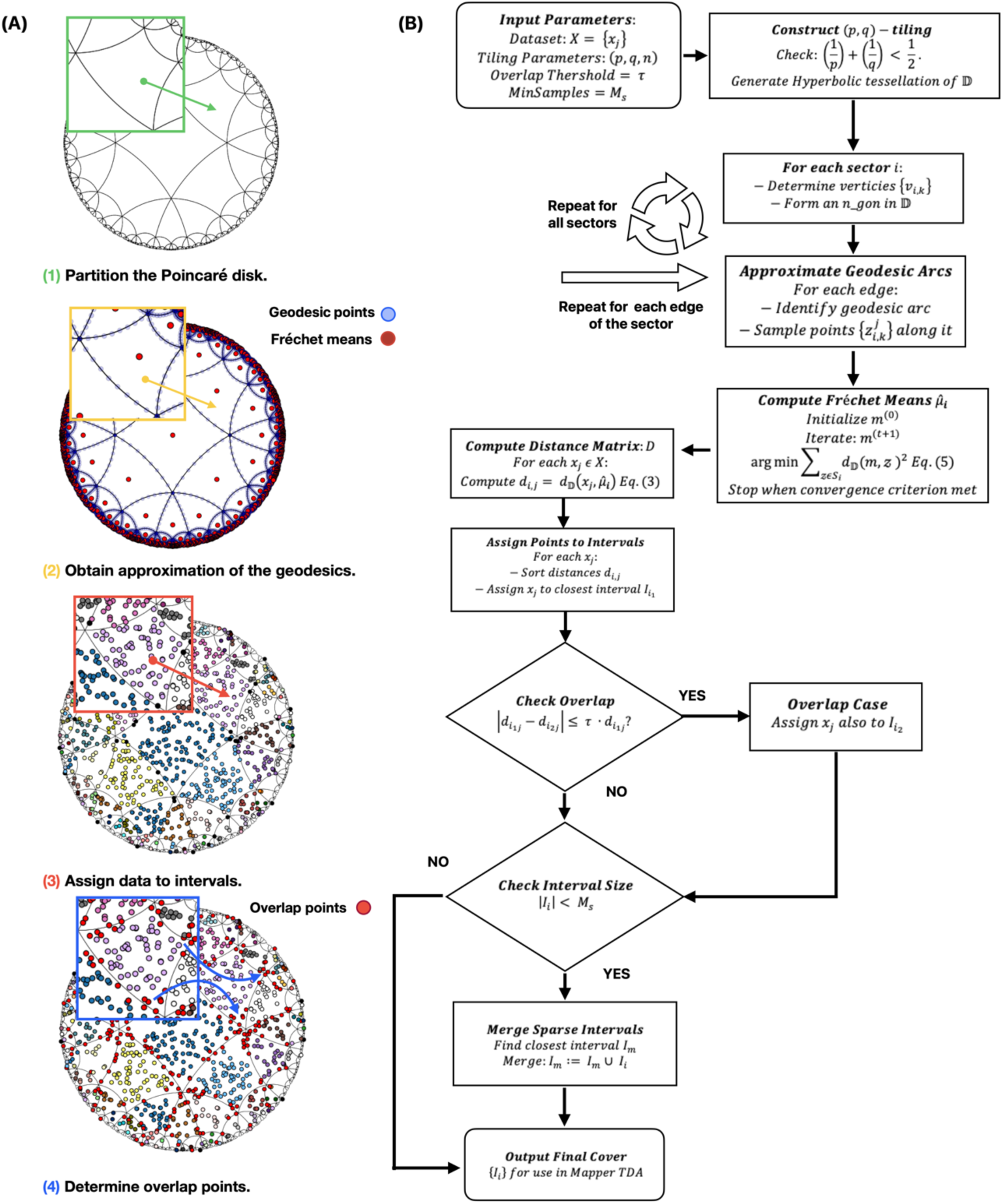
Hyperbolic space partition used for Mapper construction. (A) Regular (p,q)-tiling of the Poincaré disk, geodesic-boundary approximation, assignment of data points to intervals based on Fréchet means, and controlled overlap between adjacent regions. (B) Algorithmic workflow from disk tessellation and Fréchet mean estimation to interval merging, yielding a curvature-consistent partition that underpins subsequent Mapper and clustering analyses.

Bridge structure reinforced this interpretation. Spd-enriched regions coincided with the narrow corridors between the early hub and the late arms, including high-betweenness edges. Both distal arms were enriched for Spm, but each was approached through a different transition corridor. The graph therefore resolves two late rosette programs that share early history but differ in the path by which they are reached.

### Nutrition, dose, and transporter genotype alter occupancy rather than topology

Nutrition modified the probability of occupying developmental regions while preserving the same global topology (Figure 2A-2). The raw dataset was nearly balanced between low-MS and full-MS conditions, but low-MS samples were relatively enriched in the early hub, whereas full-MS samples were more frequent in distal regions. Nutrient availability therefore shifts residence toward or away from late compact morphologies rather than generating a second manifold.

Dose effects were weaker and more graded than the effect of polyamine identity (Figure 2A-4). Unprimed and lowest-dose samples were concentrated toward interior regions of the graph, whereas higher non-zero levels appeared more often along the periphery and distal Spm-rich arms. This pattern supports dose-dependent modulation of how far plants penetrate late morphological states within the observed time window, although we interpret the effect as secondary to priming identity and nutrition.

**Figure 4.**
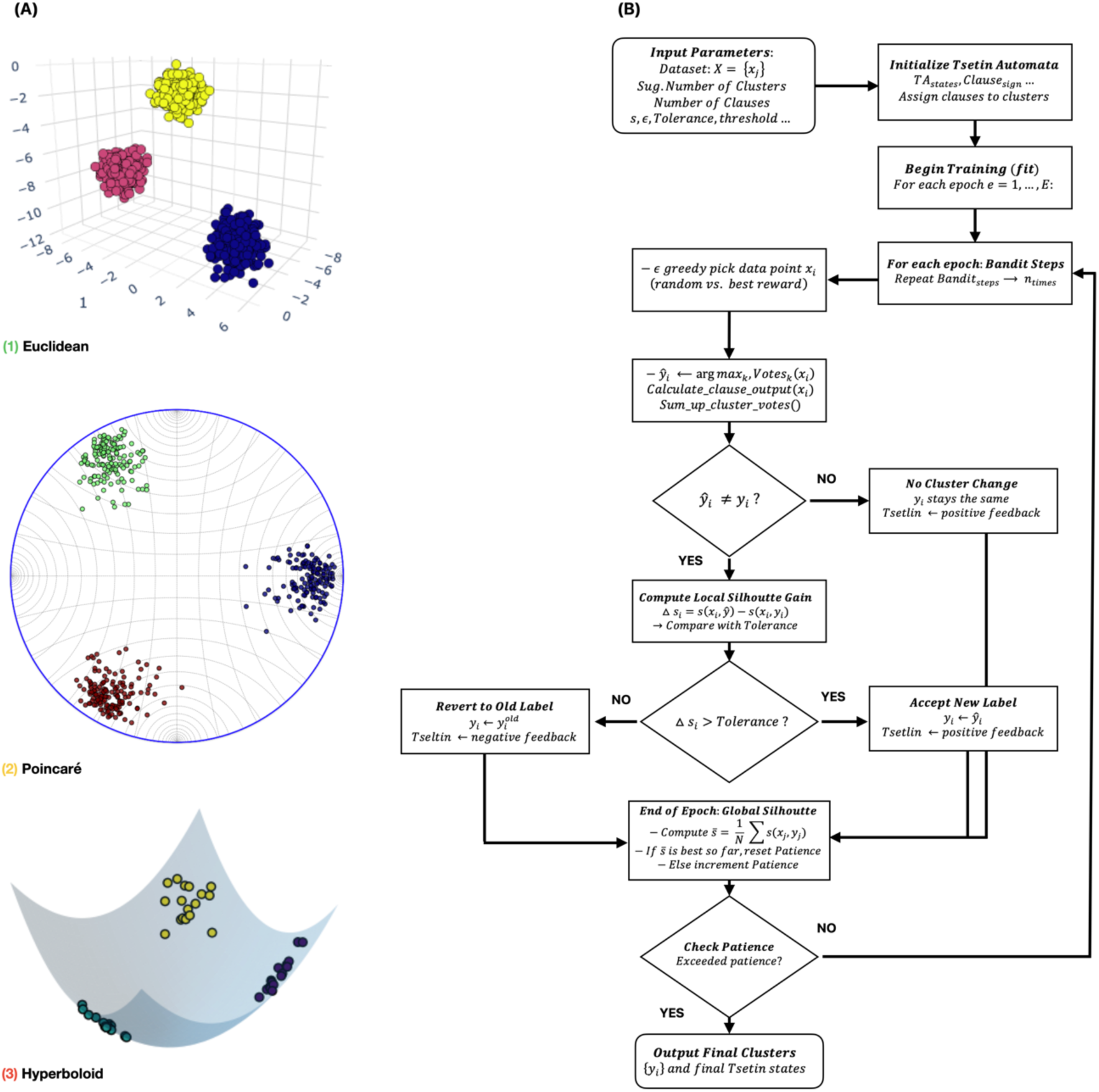
Logic-driven clustering across latent geometries. (A) Example outcomes in Euclidean, Poincaré disk, and hyperboloid representations. (B) Workflow showing clause-based voting, silhouette-guided feedback, and multi-armed-bandit refinement. The framework yields cohesive, interpretable clusters across distinct geometric representations of the latent space.

Genotype had the smallest effect on manifold geometry (Figure 2A-5). Col-0 and all five *atput* mutants occupied the same connected structure, including the early hub, transition regions, and late arms. Local enrichments were visible, with Col-0 slightly overrepresented in one distal lobe, *atput1/3/5* more frequent around early-to-mid connectors, and *atput4* distributed broadly across the graph. These differences are consistent with altered dwell times or transition probabilities within shared states, rather than with genotype-specific morphotypes.

### Manifold straightening reveals an early steering window

To determine whether the Mapper arms reflected persistent branching or transient lateral deviations, we straightened the graph-derived manifold into a ribbon coordinate system (Figure 2B-1 to B-3). A minimum-spanning-tree backbone was refined into a medial centerline; each plant was then described by normalized pseudotime s and signed transverse displacement y. The resulting ribbon was narrow for most of the trajectory and widened primarily during early pseudotime, indicating that treatment-associated variation is strongest near entry into the manifold and subsequently reconverges.

Group curves in the straightened frame localized the major effects. Put and Spm deviated toward opposite shoulders during the early widened interval, Spd and unprimed controls remained closer to the centerline, and genotype-specific curves largely overlapped. Nutrition produced a modest early separation between low-MS and full-MS samples before convergence. Thus, the ribbon representation condenses the Mapper graph into a quantitative model: early transverse offset captures the initial polyamine/nutrient steering event, whereas distal occupancy captures entry into late SPM-associated rosette programs.

Together, the graph and ribbon analyses support a shared-manifold model of rosette morphodynamics. Nutrition, polyamines, dose, and lines do not create independent morphological spaces; they redistribute plants along a common developmental geometry. This result also defines a practical scoring strategy for future screens: new treatments can be compared by their early lateral displacement and their probability of occupying distal late-state regions (Figure 2 and Figure S1).

## Discussion

This study reframes image-based plant phenotyping from trait extraction to developmental geometry. Across 138,223 Arabidopsis rosette images, the dominant organization was a shared manifold rather than a collection of treatment-specific clusters. This distinction matters because phenomics often conflates two questions: what morphologies exist in the dataset and how experimental factors change occupancy of those morphologies. In this experiment, the accessible morphological space is broadly shared; treatment and genotype act mainly by shifting residence, transition, and terminal occupancy.

At the biological level, the map positions polyamines as time-localized modulators of rosette architecture. Put enrichment in the early hub links putrescine with early expansion states. Spd enrichment in connector regions suggests a role in transition between early and late configurations. Spm enrichment at both distal arms links spermine with late, compact canopy architecture. These assignments represent image-derived morphodynamic signatures rather than direct biochemical causality; targeted metabolite profiling, temporal rescue experiments, and transporter complementation will be needed to test mechanism.

Nutrient effects clarify the environmental component of this model. Full-MS conditions increased occupancy of distal regions, whereas low-MS conditions favored early residence. This pattern aligns with the expectation that nutrient availability affects developmental progression, but the topology indicates that nutritional changes alter access to shared states rather than create new ones. The same logic applies to concentration: dose modulates the depth of progression into late regions, especially in Spm-rich areas, but does not reorganize the manifold.

*Atput* mutant lines provide a genetic test of the shared-manifold hypothesis. If individual transporters created unique morphogenetic programs, mutant lines would be expected to form isolated graph components or genotype-specific branches. Instead, all lines occupied the same connected structure. Local enrichments therefore more plausibly reflect altered rates of entry, retention, or exit from shared states. This interpretation is consistent with functional redundancy among transporter-family members and with the idea that polyamine transport tunes progression rather than specifies an entirely new architecture.

Methodologically, the study demonstrates why geometry-aware analysis is valuable for large phenotyping screens. Standard scalar traits summarize size or color but discard the arrangement of samples in morphological space. Conventional projections can visualize this space but often encourage cluster narratives that are sensitive to the choice of projection parameters.[13,14] By combining self-supervised visual features, hyperbolic embedding, Mapper reconstruction, and ribbon straightening, the workflow retains both global topology and local deviations. The workflow produces not only a visualization but also measurable quantities that can be tested across experiments.

Two aspects of the current analysis are especially useful for future work. First, early transverse offset provides a compact signature of treatment steering before trajectories converge. Second, distal occupancy quantifies how strongly a condition drives plants into late rosette programs within a fixed time window. These metrics convert a high-dimensional image manifold into scalar phenotypes while preserving the context of the manifold from which they were derived.

Several limitations constrain interpretation. The analysis is image-based and therefore cannot, by itself, identify the molecular state underlying a node. Area-derived pseudotime is an auxiliary descriptor, not a replacement for chronological age or lineage-resolved growth rate. Mapper topology depends on embedding quality, cover design, overlap, and clustering parameters, and metadata-informed hyperparameter selection requires transparent reporting. Because Mapper cover sets overlap, node-level enrichments are best interpreted as occupancy tendencies rather than mutually exclusive states.

The broader implication is that developmental phenomics can move beyond endpoint comparisons toward atlases of possible form. In such atlases, perturbations are described by how they bend, delay, or stabilize a trajectory through shared space. Applied to polyamine biology, this view suggests that Put, Spd, and Spm are not simply growth promoters or inhibitors; they bias different phases of a common morphodynamic program. Applied more broadly, the same framework could support comparative screens across species, organs, stress regimes, and genetic backgrounds, if imaging, metadata, validation, and mechanistic follow-up are reported with sufficient precision.[11,22]

## Experimental Section

### Plant material and experimental setup

*Arabidopsis thaliana* ecotype Columbia-0 (Col-0) and T-DNA insertion lines for members of the Arabidopsis polyamine uptake transporter family [atput1-1 (GABI_890c10), atput2-1 (SALK_119707c), atput3-1 (SALK_206472c), atput4-1 (SAIL_1275_c06), and atput5-1 (SALK_007135)] were obtained from the Salk Institute Genomic Analysis Laboratory (signal.salk.edu/cgi-bin/tdnaexpress). Homozygous lines were identified by PCR, and loss of the corresponding transcript was assessed by qPCR (Supporting Information).

Seeds were surface-sterilized as described previously.[23] They were sown on square plates (120 mm × 120 mm; P-Lab, Ref. 212358.2) containing half-strength Murashige and Skoog (MS) medium (Phytotechlab M519), pH 5.7, supplemented with 0.6% Phytagel.[24] Priming media contained putrescine (Put), spermidine (Spd), or spermine (Spm) at 0.1, 1, or 10 mM, or no priming agent as a control. Plates were stratified for 3 d at 4 °C in darkness and then transferred vertically to a growth chamber under long-day conditions (22 °C, 16 h/8 h light/dark photoperiod, 120 µmol photons m⁻² s⁻¹ PAR).

Three-day-old seedlings of comparable size were transferred under sterile conditions to 48-well plates containing full-strength (1×) or low-strength (0.5×) MS medium, with one seedling per well. Plates were sealed with perforated transparent foil to allow gas exchange and transferred to the OloPhen platform, a PlantScreen^TM^ XYZ system (Photon Systems Instruments, Drasov, Czech Republic) installed in a controlled growth chamber. The chamber was maintained under a long-day regime (22/20 °C day/night, 16 h/8 h light/dark photoperiod, 120 µmol photons m⁻² s⁻¹ PAR, 60% relative humidity), following the established OloPhen protocol.[23]

### Data preprocessing

*Arabidopsis thaliana* seedlings were imaged with top-view RGB cameras in 48-well plates. Plate-level images were processed by a custom detection and cropping workflow, adapted from previous high-throughput plate-phenotyping work, to isolate individual rosettes (Figure S3).[25] A trained object-detection network localized plant centers; detections were paired with reference grid positions via Hungarian assignment; and non-maximum suppression removed redundant bounding boxes. The final dataset contained 138,223 single-plant images annotated with imaging round, plant index, genotype, priming identity, concentration, and medium.

We upscaled the cropped images fourfold using a super-resolution model prior to feature extraction (Figure S4).[26] This step reduced pixelation and improved leaf-edge contrast while preserving the original morphology. We excluded images that could not be confidently matched to a grid position before embedding.

We used metadata for downstream aggregation and interpretation, not to define the initial visual embedding. For graph-level summaries, we interpreted overlapping Mapper memberships as node occupancy rather than as mutually exclusive sample classes.

### Latent space representation

To derive a compact representation of rosette morphology, we embedded all cropped images using contrastive representation learning. A pre-trained DINOv2 vision-transformer backbone processed each image and generated a 368-dimensional feature vector that captured shape, texture, and color information relevant to rosette architecture (Figure S5).[12]

To adapt these features to developmental geometry, we attached a Poincaré head that projected the Euclidean feature vectors into a two-dimensional hyperbolic disk. Training used a SimCLR-like contrastive objective to pull together augmented views of the same plant while separating representations from different images.[27]

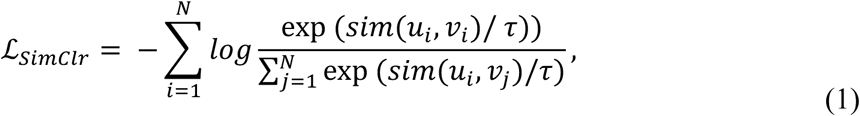

Where *sim*(*u*, *v*) denotes the similarity (e.g., cosine similarity) between two embeddings *u* and *v*, *τ* is the temperature parameter and *N* is the number of positive pairs (i.e., pairs of views created from augmentations of the same original image).

The Poincaré ball is a conformal model of hyperbolic space: it preserves angles in the tangent space while distances are governed by the hyperbolic metric rather than by the ambient Euclidean metric. This geometry is suitable for data with hierarchical or branching organization, including developmental progressions.

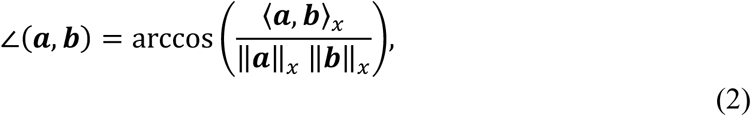

Where 〈·,·〉_+_ and ‖·‖_+_ denote the inner product and norm under the Poincaré metric at ***x***. The distance between two points ***x***, ***y*** *ε* B^#^ is defined by:

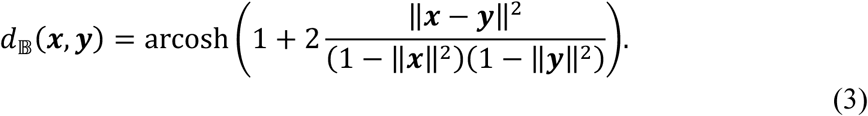

which preserves angular relationships while accommodating curvature. The Euclidean embeddings *e* are mapped into the hyperbolic disk through a curvature-controlled transformation

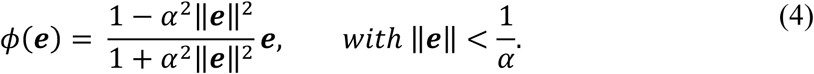

Where ***e*** is the Euclidean embedding, and *α* is a learnable (or fixed) curvature-related parameter. This formula ensures that the norm of *φ*(***e***) remains strictly less than 1, thus embedding the Euclidean data within B^#^. By incorporating this Poincaré transformation, we adapted the hybrid approach to capitalize on the geometric advantages of hyperbolic space.

The hybrid DINOv2-Poincaré representation therefore captures complementary scales of morphology: local visual similarity in the backbone features and broader developmental hierarchy in the hyperbolic embedding. The resulting two-dimensional coordinates served as the geometric substrate for Mapper graph construction and topology-aware analysis.[16–20]

For complementary validation, we reduced the 368-dimensional DINO embeddings to 10 principal-component dimensions and then applied UMAP to obtain a three-dimensional representation. This analysis verified the consistency of the global manifold geometry observed in the main Poincaré space (Figure 2B-2).[14,15] The comparison confirmed that the hyperbolic projection preserved the same temporal ordering and treatment clustering while offering clearer separation of transitional states.

This representation stage converts raw pixel information into a geometry-aware embedding that links modern representation learning with interpretable phenotyping.

### Hyperbolic Space Partitioning

After obtaining the two-dimensional embeddings in the Poincaré disk, we introduced a hyperbolic-space partition that segmented the latent manifold while preserving its intrinsic curvature. Conventional Euclidean approaches, such as cubical or grid-based covers, assume flat geometry and can distort relationships near curved boundaries. Our method instead constructed a curvature-aware tessellation that more faithfully represented complex biological data.

We generated a regular (p,q)-tiling of the Poincaré disk, where p denotes the number of polygon sides and q denotes the number of polygons meeting at each vertex. The admissible configuration satisfies (p - 2)(q - 2) > 4, yielding a hyperbolic tessellation. Geodesic edges were discretized by sampling points along circular arcs orthogonal to the unit boundary.

For every sector *i*, we calculate the Fréchet mean 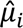 of its sampled boundary points *S_i_* by minimizing the sum of squared hyperbolic distances

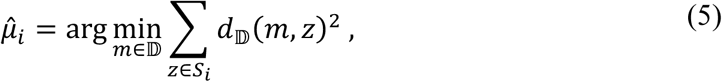

where *d*_D_(*x*, *y*) is the hyperbolic distance defined in Eq. (3). Each mean 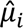 *s*erves as a geometric center representing its local neighborhood. We then computed a distance matrix *D* = [*d_ij_*], where 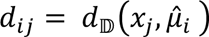 measures the distance between every data point *x*_’_ ∈ D and each sector center.

We assigned data points to cover sets according to their hyperbolic distance from sector centers. For each point, the nearest sector defined the primary interval; the second-nearest sector defined a secondary interval when its distance fell within the optimized overlap threshold. We merged sparse intervals with the nearest populated interval to preserve continuity and avoid biologically uninformative singleton regions.

These overlapping intervals formed a curvature-aware cover of the dataset. We used this cover for Mapper graph construction and clustering, ensuring that partitioning respected the geometry of the Poincaré disk rather than imposing a flat Euclidean grid (Figure 3).

### Logic-Driven Clustering in Curved Space

After assigning embeddings to hyperbolic intervals, we grouped plants within each interval using a logic-driven clustering framework. Because standard Euclidean clustering can distort distances in curved embeddings, we used a Tsetlin-machine-inspired procedure that represented samples through binary, curvature-aware distance codes and refined assignments by silhouette-guided feedback.[28]

For Euclidean features, we min-max normalized inputs and discretized them by threshold encoding. For hyperbolic embeddings, we computed distances from each point to five fixed landmarks in the Poincaré disk: one at the origin and four at a common boundary radius. We then thresholded these distances to form binary vectors.

Each Tsetlin clause comprised automata controlling the inclusion or exclusion of binary literals. Clauses voted for candidate cluster membership, and the procedure assigned each sample to the cluster that received the strongest positive vote after silhouette-guided refinement.

We optimized cluster assignment using an epsilon-greedy multi-armed bandit procedure.[29] The optimizer accepted proposed reassignments when they improved local silhouette beyond a predefined tolerance; otherwise, it reverted the assignments. We monitored global silhouette after each epoch and stopped optimization when improvement plateaued.

During bandit optimization, clauses that improved local silhouette were reinforced, whereas clauses that reduced cohesion were weakened. This feedback updated the logical representation while preserving the geometry-aware distance encoding.

The final clusters emerged from rule-based voting combined with geometry-aware silhouette feedback. This design preserved interpretability while allowing the same framework to operate on Euclidean, Poincaré, and hyperboloid representations (Figure 4).

### Area-Derived Pseudotime

To obtain an independent descriptor-based developmental score, we reused the trained visual backbone with a lightweight segmentation head. We fine-tuned the head on 3,500 manually annotated rosette masks and applied it to all images. For each plant, we defined rosette area as the number of foreground pixels in the largest rosette component after debris removal.

We computed area-derived pseudotime from the log-transformed foreground area of each plant. We ranked rosettes by area from smallest to largest and linearly scaled the ranks to a 0-1 interval, with values near 0 representing the smallest rosettes and values near 1 representing the largest rosettes. We obtained early, mid, and late summaries by splitting this global pseudotime distribution into tertiles (Figure S1).

### Multiobjective Hyperparameter Optimization

We selected hyperparameters with a multiobjective NSGA-II sampler that balanced local clustering quality and global topological coherence.[30] We checked local partitions with silhouette-type cohesion and Davies-Bouldin separation diagnostics.[31] Because the objective functions included metadata, we interpret this stage as metadata-informed model selection rather than as a fully unsupervised training step. We subsequently interpreted the selected graph by projecting time, line, concentration, nutrition, and priming metadata onto the fixed topology.

We selected the final model after 100 optimization trials. The hyperbolic cover used an overlap threshold of 0.253, a minimum interval size of 64 samples, a regular four-sided, ten-at-a-vertex tiling, and four tiling layers. The logic-driven clustering stage used 312 clauses, 171 automaton states, a specificity setting of 3.9, a voting threshold of 15, 300 bandit-refinement steps, a negative-silhouette tolerance of 0.01, and an exploration probability of 0.3.

These jointly optimized parameters balanced local cohesion, global topological integrity, and computational efficiency, yielding the final configuration used throughout the study.

### Contrastive Learning Configuration

For contrastive learning, we normalized the complete image set using channel-wise statistics from the training data. We generated two augmented views per image using random resized cropping, horizontal flipping, color jitter, and Gaussian blur.

Training used the Adam optimizer for the DINOv2 backbone and Riemannian Adam for the Poincaré head to maintain curvature-consistent updates.[32] The head comprised Poincaré-linear layers, Poincaré batch normalization, and a final Poincaré-linear output layer producing two-dimensional embeddings within the hyperbolic disk.

Optimization used an InfoNCE-type contrastive loss with temperature parameter τ, following the SimCLR-style training configuration described above.[27] Training proceeded for up to 200 epochs and stopped early when validation performance plateaued.

## Acknowledgements

This work was funded by the European Regional Development Fund (ERDF) Programme Johannes Amos Comenius (CZ.02.01.01/00/23_021/0008909) and Ciencia Básica y de Frontera (CBF-2025-I-3650) funded by Secretaría de Ciencia, Humanidades, Tecnología e Innovación (SECIHTI).

## Conflict of Interest

The authors declare no conflict of interest.

## Author Contributions

**Jan Zdražil:** Conceptualization, Investigation, Formal analysis, Writing original draft, Visualization. **LingPing Kong:** Investigation, Formal analysis. **Emmanuel Flores-Hernández:** Investigation, Formal analysis. **Margarita Rodríguez-Kessler:** Investigation, Formal analysis, Resources, Funding acquisition. **Pavel Klimeš:** Data curation, Investigation. **Lukáš Spíchal:** Conceptualization, Writing – review & editing, Resources, Funding acquisition. **Nuria De Diego:** Conceptualization, Data curation, Formal analysis, Writing – original draft, Writing review & editing, Resources, Funding acquisition. **Václav Snášel:** Conceptualization, Investigation, Writing review & editing, Resources, Funding acquisition.

## Data Availability Statement

The source code used for image preprocessing, embedding, and manifold analysis will be deposited in a public repository or provided as Supporting Information prior to submission. The raw and processed image datasets generated and analyzed during the current study are available from the corresponding author on reasonable request, subject to institutional data-transfer requirements.

## Table of Contents Text

High-throughput images of *Arabidopsis* rosettes are converted into a hyperbolic topological atlas. Across nutrient regimes, polyamine treatments, doses, and *Atput* transporter mutants, plants occupy a shared developmental manifold rather than separate morphologies. Putrescine, spermidine, and spermine bias early, transition, and late states, enabling compact metrics for morphodynamic phenotyping.

## Supporting Information

**Figure S1.**
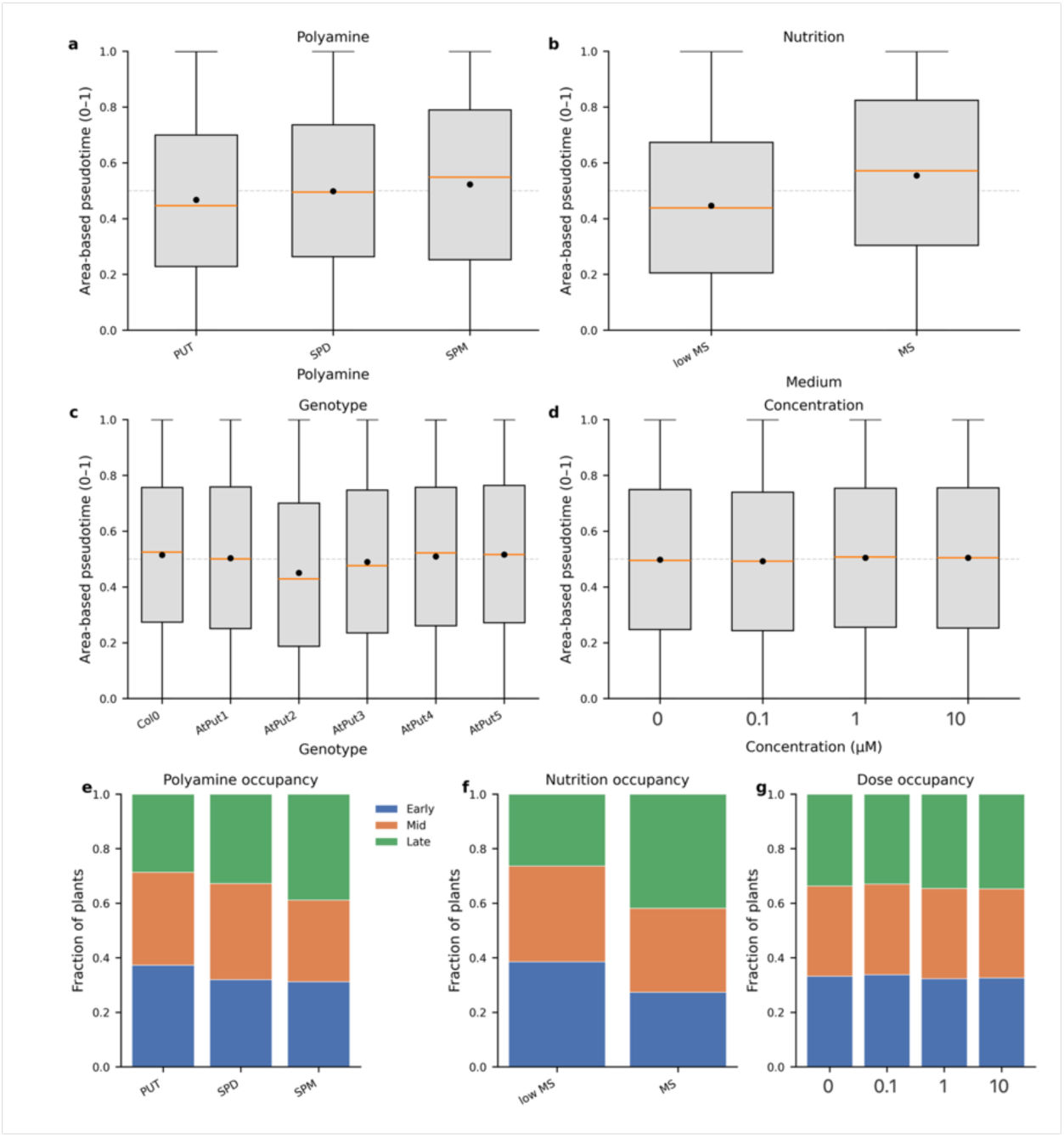
Area-derived pseudotime summaries by polyamine identity, nutrition, genotype, and dose. (a-d) Each point represents one rosette image; black dots mark group means and the dashed grey line marks the global mean across approximately 138,000 plants. Put-primed rosettes shift toward earlier pseudotime, Spm-primed rosettes shift toward later states, and Spd remains near the global mean. Low-MS medium biases plants toward earlier values, whereas full MS shifts them later; the five Atput mutant lines and Col-0 span a narrow shared range. Dose effects are detectable but modest compared with polyamine identity or nutrition. (e-g) Stacked bars show the fraction of plants in early, mid, and late tertiles of area-based pseudotime for polyamine identity, nutrition, and dose. These descriptor-based summaries support the conclusion that treatments modulate residence along a shared morphodynamic trajectory rather than generating distinct developmental routes.

**Figure S2.**
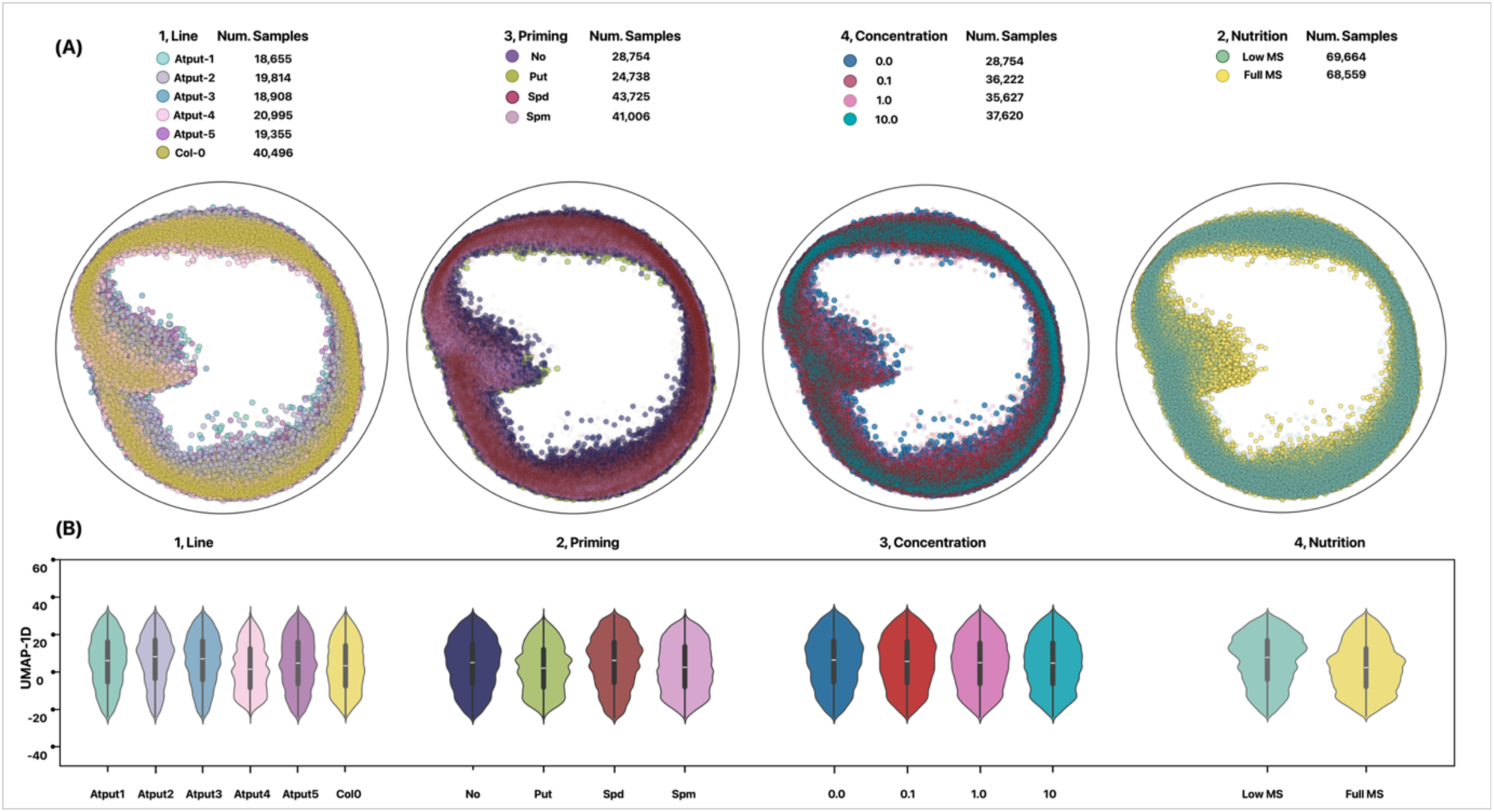
Two-dimensional Poincaré embeddings obtained from the filter function, visualized by line, nutrition, priming, and concentration. The substantial overlap and hierarchical structure across conditions highlight the complexity of the dataset. Violin plots after further reduction from two dimensions to one dimension by UMAP show strong overlap, motivating the use of topology-aware analyses rather than simple projection-based summaries.

**Figure S3.**
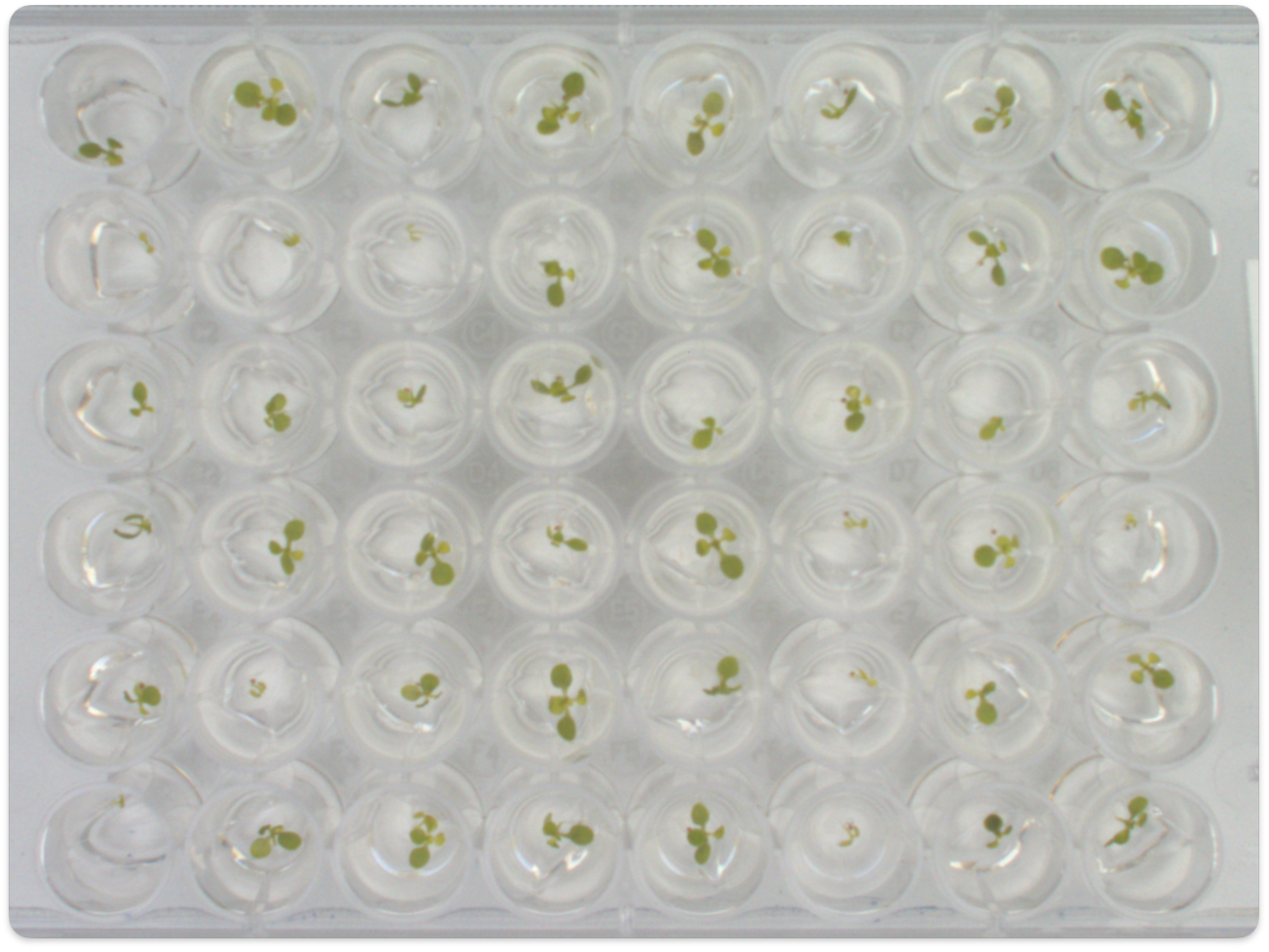
Representative RGB image of the 48-plant plate configuration used for data acquisition. Each well contains a single Arabidopsis thaliana seedling imaged under controlled illumination and camera geometry, forming the raw dataset used in downstream analyses.

**Figure S4.**
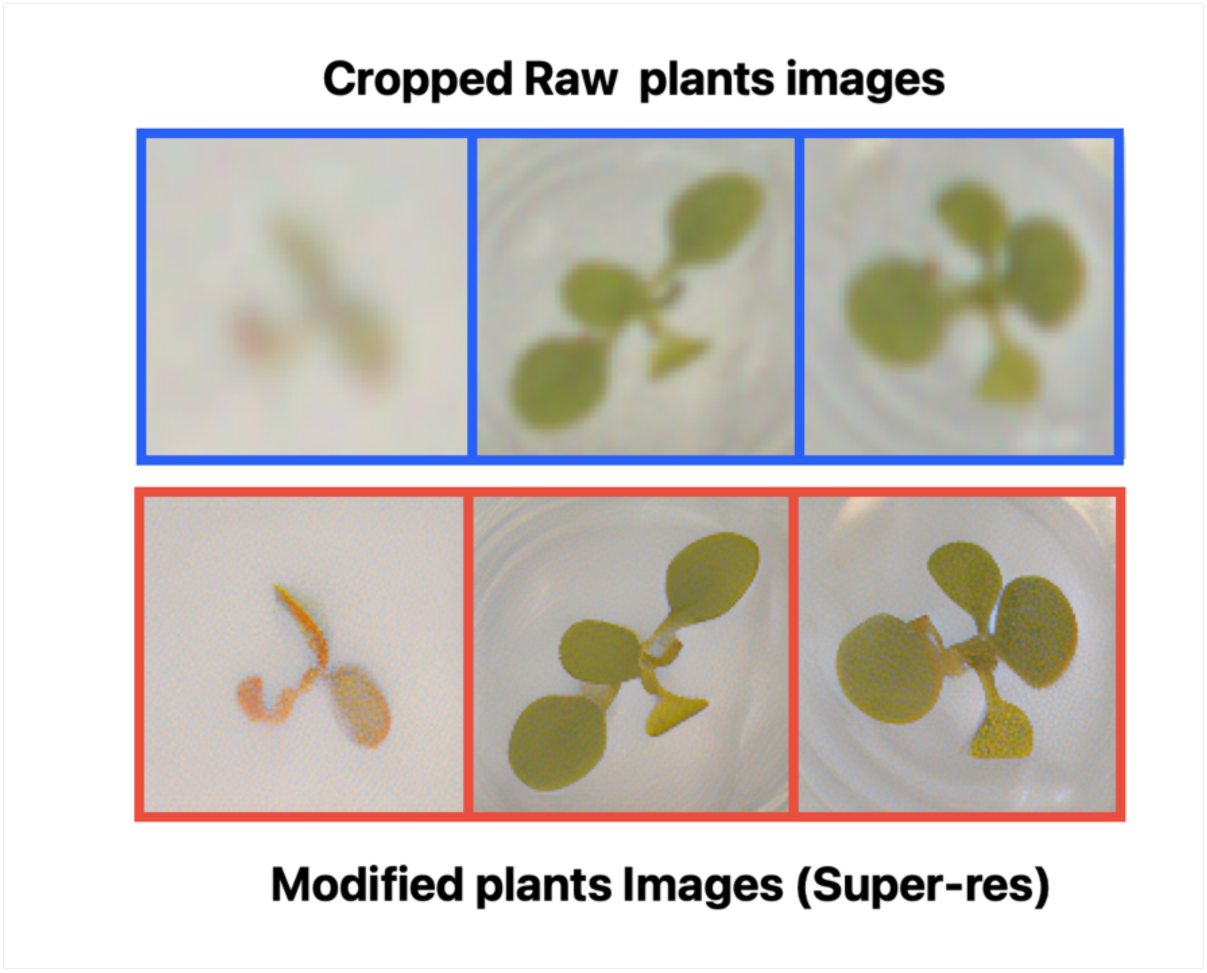
Original cropped plant images extracted from the detection pipeline and corresponding outputs after super-resolution processing. The enhanced images show sharper leaf edges and improved image fidelity while preserving rosette morphology.

**Figure S5.**
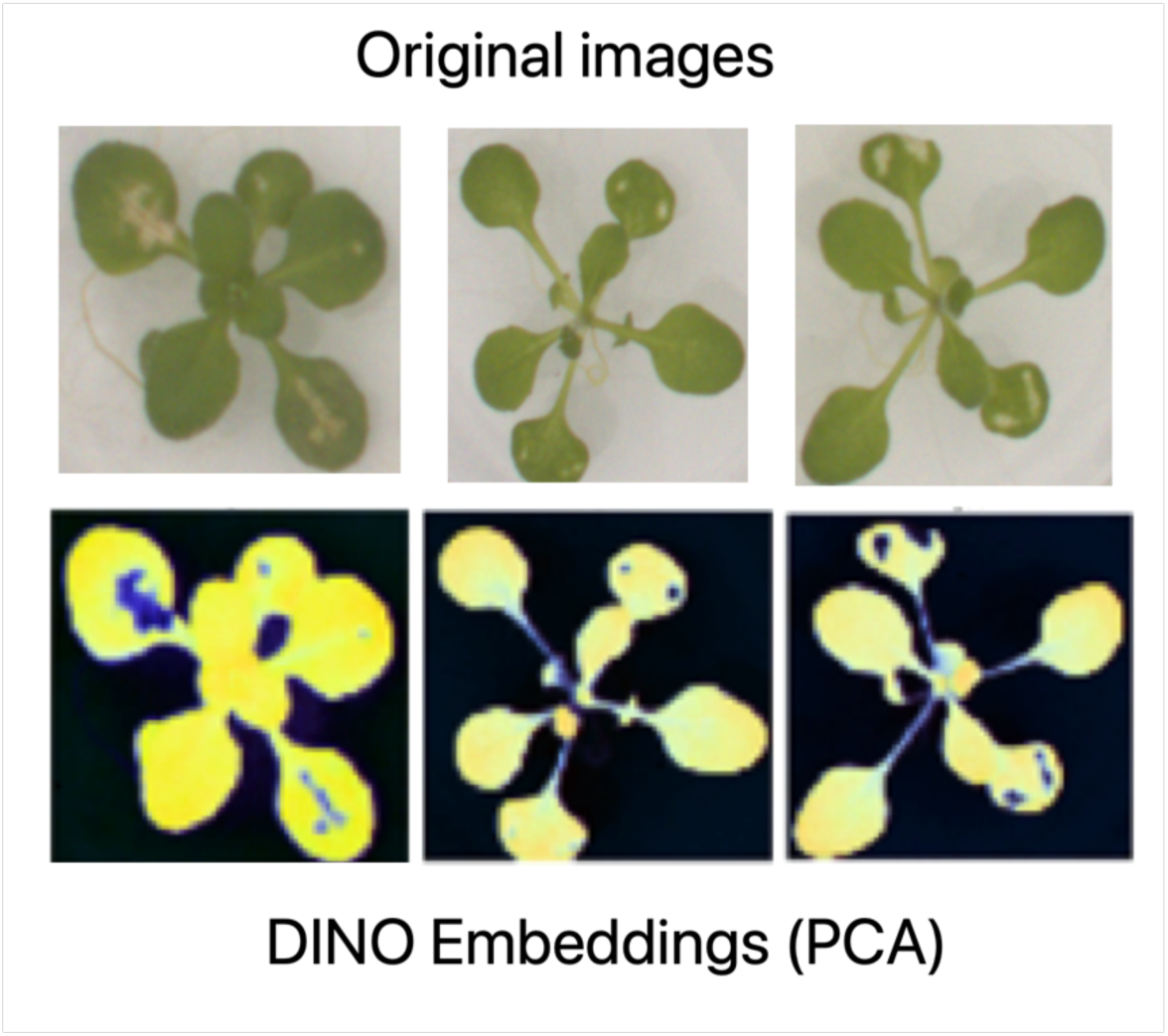
Representative cropped plants used as input examples and their corresponding high-dimensional DINOv2 embeddings visualized by PCA projection.

## Notes

### Competing Interest Statement

The authors have declared no competing interest.

## References

[1] F. Fiorani, U. Schurr, Annu. Rev. Plant Biol. 2013, 64, 267.

[2] A. Zavafer, H. Bates, C. Mancilla, P. J. Ralph, Trends Plant Sci. 2023, 28, 1004.

[3] W. Yang, H. Feng, X. Zhang, J. Zhang, J. H. Doonan, W. D. Batchelor, L. Xiong, J. Yan, Mol. Plant 2020, 13, 187.

[4] K. M. Murphy, E. Ludwig, J. Gutierrez, M. A. Gehan, Annu. Rev. Plant Biol. 2024, 75, 771.

[5] G. D. Tan, U. Chaudhuri, S. Varela, N. Ahuja, A. D. B. Leakey, J. Exp. Bot. 2024, 75, 6683.

[6] L. Yu, H. Sussman, O. Khmelnitsky, M. R. Ishka, A. Srinivasan, A. D. L. Nelson, M. M. Julkowska, Plant Physiol. 2024, 196, 810.

[7] M. C. Heuermann, D. Knoch, A. Junker, T. Altmann, Nat. Commun. 2023, 14, 5783.

[8] D. Hobby, H. Tong, M. Heuermann, A. J. Mbebi, R. A. E. Laitinen, M. Dell’Acqua, T. Altmann, Z. Nikoloski, Nat. Plants 2025, 11, 1018.

[9] A. L. Harfouche, F. Nakhle, A. H. Harfouche, O. G. Sardella, E. Dart, D. Jacobson, Trends Plant Sci. 2023, 28, 154.

[10] X. Li, C. Xie, L. Cheng, H. Tong, R. Bock, Q. Qian, W. Zhou, Trends Biotechnol. 2025, 43, 2479.

[11] F. Chen, I. Stogiannidis, A. Wood, D. Bueno, D. Williams, F. Macfarlane, B. D. Grieve, D. Wells, J. A. Atkinson, M. J. Hawkesford, S. A. Rolfe, T. Lawson, T. Pridmore, S. A. Tsaftaris, M. V. Giuffrida, Nat. Commun. 2026, DOI 10.1038/s41467-026-71090-y.

[12] M. Oquab, T. Darcet, T. Moutakanni, H. Vo, M. Szafraniec, V. Khalidov, P. Fernandez, D. Haziza, F. Massa, A. El-Nouby, M. Assran, N. Ballas, W. Galuba, R. Howes, P.-Y. Huang, S.-W. Li, I. Misra, M. Rabbat, V. Sharma, G. Synnaeve, H. Xu, H. Jégou, J. Mairal, P. Labatut, A. Joulin, P. Bojanowski, Trans. Mach. Learn. Res. 2024.

[13] M. Meilă, H. Zhang, Annu. Rev. Stat. Appl. 2024, 11, 393.

[14] J. Healy, L. McInnes, Nat. Rev. Methods Primers 2024, 4, 82.

[15] M. Greenacre, P. J. F. Groenen, T. Hastie, A. I. D’Enza, A. Markos, E. Tuzhilina, Nat. Rev. Methods Primers 2022, 2, 100.

[16] A. Klimovskaia, D. Lopez-Paz, L. Bottou, M. Nickel, Nat. Commun. 2020, 11, 2966.

[17] Y. Xu, Z. Zang, B. Hu, J. Xia, S. Z. Li, Brief. Bioinform. 2025, 26, bbae687.

[18] P. Mettes, M. Ghadimi Atigh, M. Keller-Ressel, J. Gu, S. Yeung, Int. J. Comput. Vis. 2024, 132, 3484.

[19] K. R. Moon, D. van Dijk, Z. Wang, S. Gigante, D. B. Burkhardt, W. S. Chen, K. Yim, A. van den Elzen, M. J. Hirn, R. R. Coifman, N. B. Ivanova, G. Wolf, S. Krishnaswamy, Nat. Biotechnol. 2019, 37, 1482.

[20] M. Saggar, O. Sporns, J. Gonzalez-Castillo, P. A. Bandettini, G. Carlsson, G. Glover, A. L. Reiss, Nat. Commun. 2018, 9, 1399.

[21] M. A. Blázquez, Annu. Rev. Plant Biol. 2024, 75, 95.

[22] J. E. Politsch, A. González-Delgado, K. Wabnik, Curr. Opin. Biotechnol. 2026, 97, 103428.

[23] N. De Diego, T. Fürst, J. F. Humplík, L. Ugena, K. Podlešáková, L. Spíchal, Front. Plant Sci. 2017, 8, 1702.

[24] T. Murashige, F. Skoog, Physiol. Plant. 1962, 15, 473.

[25] J. Zdražil, L. Kong, P. Klimeš, F. I. Jasso-Robles, I. Saiz-Fernández, F. Güder, L. Spíchal, V. Snášel, N. De Diego, Comput. Electron. Agric. 2025, 235, 110390.

[26] R. Rombach, A. Blattmann, D. Lorenz, P. Esser, B. Ommer, in Proc. IEEE/CVF Conf. Comput. Vis. Pattern Recognit. (CVPR), IEEE, Piscataway, NJ 2022, pp. 10684–10695.

[27] T. Chen, S. Kornblith, M. Norouzi, G. Hinton, in Proc. Int. Conf. Mach. Learn., PMLR 2020, pp. 1597–1607.

[28] L. Jiao, X. Zhang, O.-C. Granmo, K. D. Abeyrathna, IEEE Trans. Pattern Anal. Mach. Intell. 2023, 45, 6072.

[29] P. Auer, N. Cesa-Bianchi, P. Fischer, Mach. Learn. 2002, 47, 235.

[30] K. Deb, A. Pratap, S. Agarwal, T. Meyarivan, IEEE Trans. Evol. Comput. 2002, 6, 182.

[31] D. L. Davies, D. W. Bouldin, IEEE Trans. Pattern Anal. Mach. Intell. 1979, PAMI-1, 224.

[32[ G. Bécigneul, O.-E. Ganea, in Proc. Int. Conf. Learn. Represent. (ICLR), 2019.

